# Big root approximation of site-scale vegetation water uptake

**DOI:** 10.1101/559237

**Authors:** Martin Bouda

## Abstract

Land surface model (LSM) predictions of soil moisture and transpiration under water-limited conditions suffer from biases due to a lack of mechanistic process description of vegetation water uptake. Here, I derive a ‘big root’ approach from the porous pipe equation for root water uptake and compare its predictions of soil moistures during the 2010 summer drought at the Wind River Crane site to two previously used Ohm’s law analogue plant hydraulic models. Structural error due to inadequate representation of root system architecture (RSA) in both Ohm’s law analogue models yields significant and predictable moisture biases. The big root model greatly reduces these as it better represents RSA effects on pressure gradients and flows within the roots. This new mechanistic model advances our understanding of vegetation water limitation at site scale with potential to improve LSM predictions of soil moisture, temperature and surface heat, water, and carbon fluxes.

## Introduction

Vegetation access to soil water can play a key role in limiting terrestrial fluxes of water [1], heat [2] and carbon dioxide [3, 4], yet it remains poorly understood at relevant scales. A lack of accurate descriptions of plant water limitation gives rise to significant prediction errors in terrestrial or land surface models (LSM), which aim to predict these surface fluxes as a bottom boundary condition on atmospheric circulation in earth system simulations [5, 6, 7]. Results from Coupled Model Intercomparison Project Phase 5 (CMIP5) simulations of the Intergovernmental Panel on Climate Change (IPCC) show overestimates of evapotranspiration (ET) and associated overland cooling in most terrestrial regions, especially under soil-moisture limited ET regimes [8], while simultaneously underestimating terrestrial drying and the emergence of those regimes [9]. CMIP5 simulations underline the critical role of plant access to variable soil moisture in mediating surface-atmosphere fluxes and vegetation dynamics [10]. This study therefore focuses on mechanistic description of vegetation access to soil moisture as a means of improving predictions of soil moisture, vegetation water uptake, and attendant fluxes in terrestrial models.

A common approach in large-scale land surface, dynamic vegetation or ecosystem dynamics models is to calculate water uptake from each soil layer in proportion to an assumed root length density fraction reduced by a stress factor, which is often a linear function of local soil moisture status [11, 12, 13, 14, 15, 16, 17, 7, 18, 19, 20, 21]. It has been shown that changing this root water uptake formulation can give rise to significant differences in predicted transpiration across models [22, 23]. As awareness of the importance of plant hydraulics for ecosystem ecology and earth system science has grown [24], it has become apparent that more mechanistic descriptions of plant hydraulics are needed in LSMs to capture vegetation behaviour during drought [25], especially since the frequency of such water-limiting events is likely to increase in the future [26].

This has given an impetus to replace the classical plant water stress formulation with a more process-based model [27, 28, 29], based on an Ohm’s law analogue for water flow in plants [30]. While such an approach works well for stems, its weakness in representing root water uptake is that it places all plant resistances between the root base and each soil layer either exclusively in parallel [30] or in series [31]. Each of these configurations is an end-member of the spectrum of possible root system architectures (RSA) in that all other RSAs combine resistances to within-plant flow between layers in series with resistances in parallel. The actual arrangment of resistances in RSA is crucial to controlling uptake from variably saturated soils [32, 33, 34, 35, 36], as it determines the rate at which water uptake dissipates water potential gradients [37, 38], which drive flow along the soil-plant-atmosphere continuum [30]. As water uptake in a given layer dissipates the potential gradient between adjacent layers, it reduces how much water can be taken up in deeper layers. The strength of this effect depends on the soil water potentials in adjacent layers and the full arrangement of resistances. Whereas the parallel resistance structure (Fig. 1a) only allows uptake in one layer to affect others via the potential at the root base, the model that places all layers in series (Fig. 1c) assumes that the plant water potential value effective in uptake and in transport are the same in a given layer, oversimplifying the effects of adjacent water potential values on each other. Neither is a suitable conceptual model for the complex cross-layer effects known to emerge in spatially explicit RSA simulations [37].

**Figure 1:**
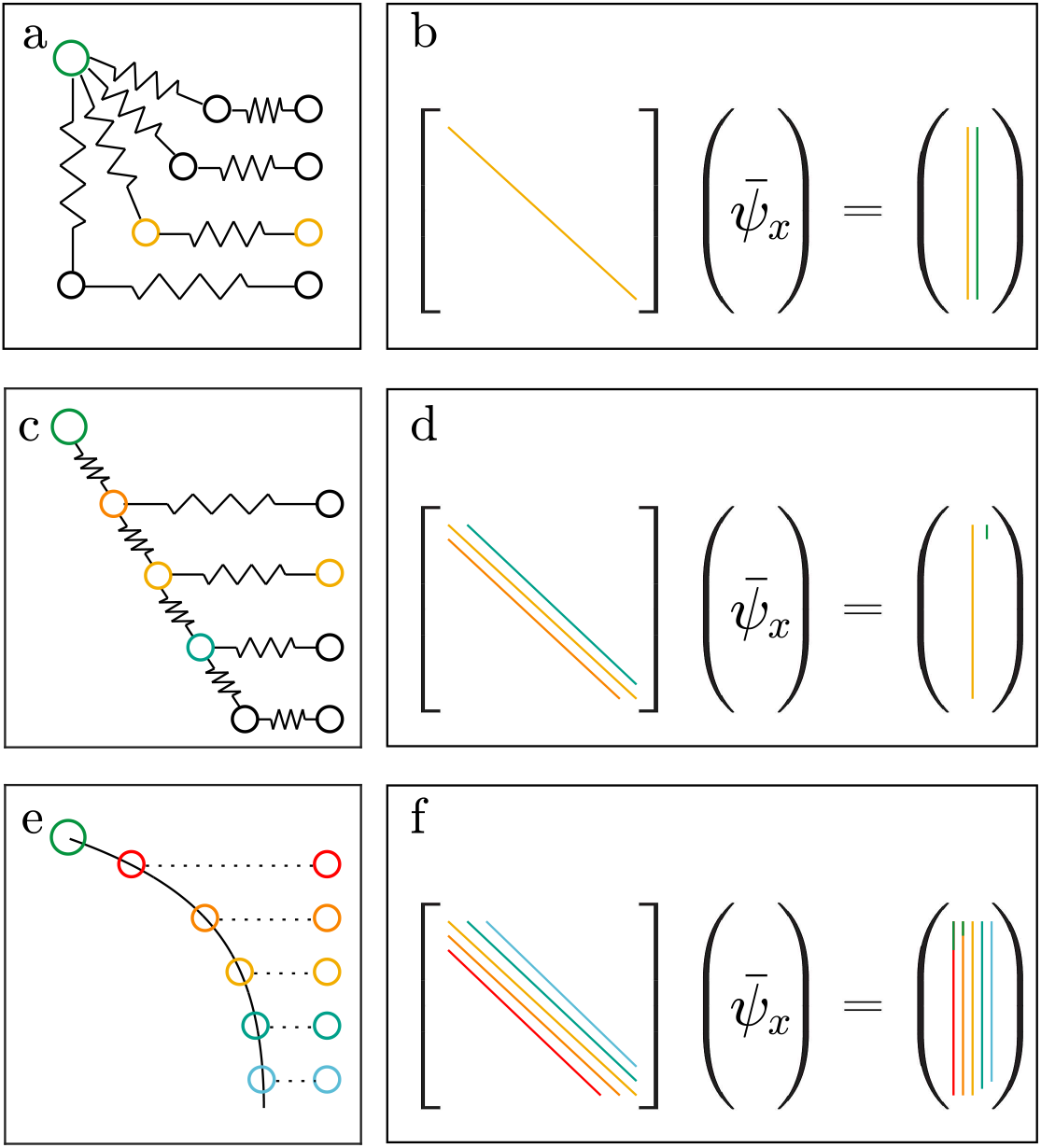
Comparison of conceptual models: Resistance diagrams for the Ohm’s law models with subsequent soil layers connected to the root base in parallel (a) and in series (c). Conceptual diagram of big root model (e) as a solution to the continuous, non-linear porous pipe equation. Circles represent water potentials at the root base (green, top left), in the soil (right-most column) and within the root (inner nodes in resistance networks in a and c, and along the solid root water potential curve in e). Sparsity patterns and composition of linear systems for layer root water potential 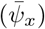 in the Ohm’s law parallel (b), series (d), and big root (f) models, colour-coded to correspond to elements in conceptual diagrams. Coefficient matrices show non-zero diagonals present in each model, with yellow on the main diagonal (layer being solved for), orange and red for layers above and dark and pale blue for layers below. Right hand side vector entries are linear combinations of the soil water potentials of individual layers, identically colour coded. Green denotes a term for the root base boundary condition. Coloured nodes in conceptual diagrams show elements involved in the solution for root water potential in yellow layer.

A conceptual alternative is provided by the ‘porous pipe’ model of root water uptake [39], which describes the continuous variation of water potential (*ψ*) along the length (*s*) of roots taking up water with the differential equation

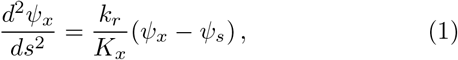

where the left-hand side term represents the dissipation of the gradient of root water potential (*ψ_x_*), which is proportional to the water potential difference between the root and the soil (*ψ_s_*), with the ratio of conductance into (*k_r_*) and along (*K_x_*) the root as the constant of proportionality. Analytical solutions to this equation have recently been extended from the case of single roots with homogeneous properties to arbitrary RSAs [40, 38]. This paves the way for solutions to this continuous model that predict the root water potential effective in water uptake in each layer 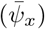 as a function of varying soil water potentials and canopy demand (Fig. 1e). Here, I show that using the porous pipe equation to find 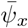 analytically under the simplifying assumption that all roots at a site can be represented with a single ‘big root’ improves vegetation water uptake predictions during a drought at a forested site, compared to the Ohm’s law models, as the resulting model better represents the effects of root system architecture on belowground pressure gradients.

## Results

### Model Form

For a single root traversing soil layers of uniform water potential, the water potential effective in water uptake in the *i*^th^ layer can be expressed as

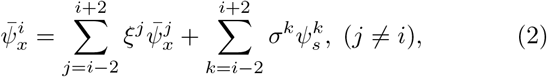

with linear coefficients *ξ* and *σ* derived analytically from root properties (hydraulic conductances *k_r_, K_x_*, and root length *S*, see Methods and Table S1). Assuming that the root systems of vegetation at a given site collectively behave as a single effective root, will be termed the ‘big root’ approach.

This ‘big root’ model differs from the two Ohm’s law models in how it that it reflects more complex interactions between water potentials across layers, effectively capturing more realistic effects of RSA on water uptake. Each of the models can be formulated as a linear system, whose solution is the vector of root water potential values for each layer, depending on soil water potentials and the stem base boundary condition. Due to their linearization of the root resistances in a specific, assumed RSA arrangement, both Ohm’s law models retain a limited set of cross-layer interactions. The layers-in-parallel Ohm’s law model (fig. 1a) assumes each layer’s root water potential value depends only on the same layer’s soil water potential and the root base boundary condition (Fig. 1b). The layers-in-series model (fig. 1c) assumes it depends instead on the root water potentials in the layers above and below, as well as the same layer’s soil water potential; only in the top layer is the root base boundary condition relevant (Fig. 1d). In the big root model (fig. 1e), by contrast, the root water potential in a given layer depends on the root and soil water potentials of two layers above and the two below; only in the two topmost layers is the root base boundary condition relevant (Fig. 1f). The big root model thus represents more complex interactions among depth layers without noticeably increasing the computational cost of the calculation (SI Numerical Results).

### Numerical Simulations

Each model was used to predict vegetation water uptake and soil moisture during the 2010 summer drought at the Wind River Crane site [41, 42, 43].

By setting root water uptake or outflow into the soil as a linear function of the difference between stem base and soil water potential in a given layer, the layers-in-parallel Ohm’s law model (Fig. 1a,b) allows for spurious soil moisture redistribution as water flows from soil layer to layer via the stem base. As a result, this mdoel had the highest overall root mean squared error (RMSE) in soil moisture predictions (10.32% summed across layers), with both dry biases in relatively wet layers and wet biases in dry layers (Fig. 2, left). In the shallow soil layer, this led to increasing nocturnal water outflow over time, even as the model predicted nocturnal excessive uptake at 50 and 100cm depths (Fig. 3a-b,g,k-l). This redistribution so depletes the 50cm layer over the first half of the simulation (Fig. 3g) that it is unable to to support observed root water uptake later on (Fig. 3h), furhter increasing spurious uptake at 100cm (Fig. 3l). The model accumulates least bias at the 30cm depth by the end of the simulation, but this is because its initial overprediction of peak uptake rates (Fig. 3c) is compensated in the second half of the simulation by spurious nocturnal water redistribution into the layer (Fig. 3d). The assumed RSA thus leads to excessive smoothing of the soil moisture profile over time, with wet biases in dry soil layers and vice versa, consistent with previous findings [29].

The RSA assumed by the layers-in-series model (Fig. 1c,d) imposes rigid water potential gradients across layers, leading to overestimation of uptake in shallow layers subjected to overly negative water potentials and underestimation of uptake at depth, where only slightly negative potentials at depth are sustained (Fig. 3). The pattern holds at the half-hourly scale, for all times and throughout the simulation period (Fig. 4 centre). As a result, while this model performs slightly better for these data overall (8.15% RMSE across all layers), it consistently yields dry biases in shallow soil layers and wet biases in deep layers (Fig 2, centre).

**Figure 2:**
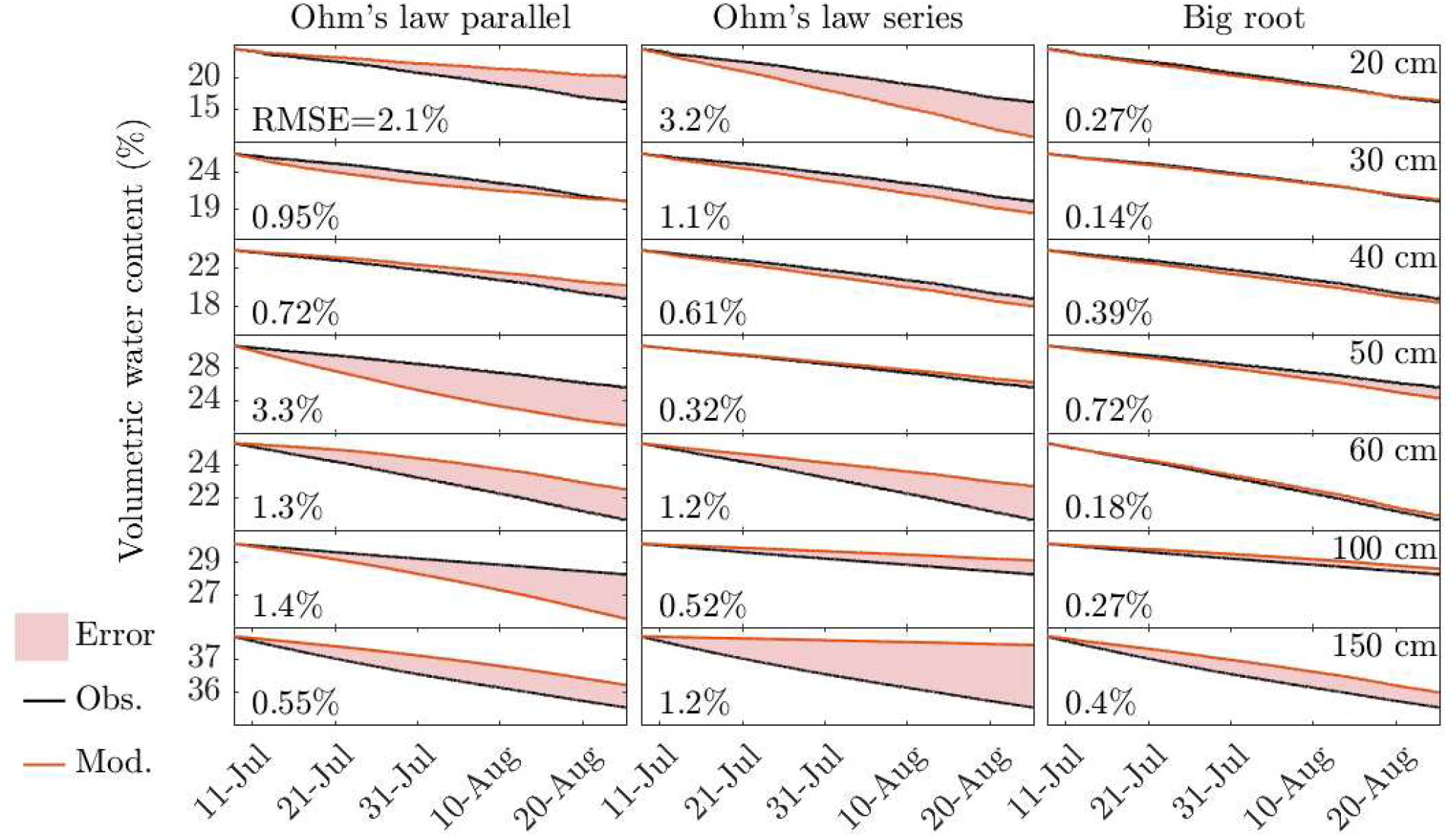
Observed (black) and predicted (red) volumetric soil moisture (%) over the entire simulation period for the Ohm’s law parallel (left), series (centre), and big root (right) models, divided by soil layers (depths at right). Prediciton errors shaded in red. Root Mean Square Error of prediction reported at left of each panel.

**Figure 3:**
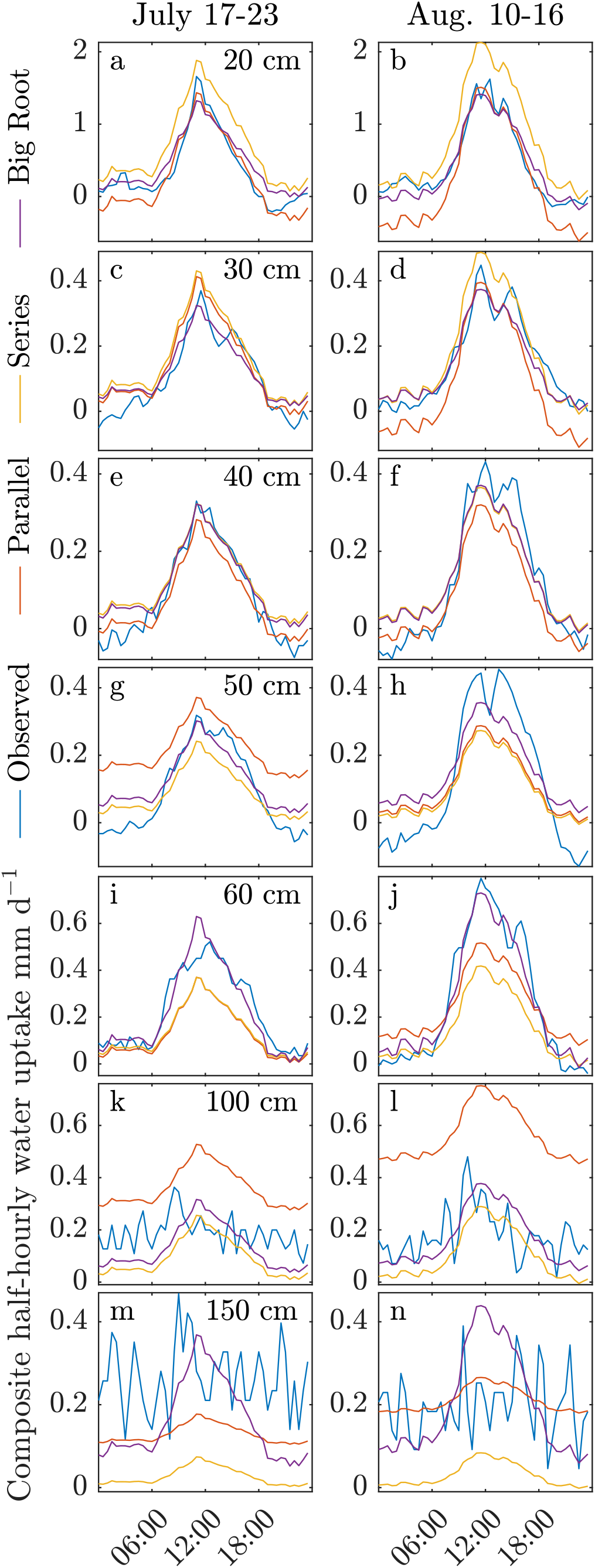
Comparison of observed and modelled half-hourly water uptake rate (mm/d) for 6-day composites of periods outside calibration dataset, displayed separately for each depth layer.

The big root model (Fig. 1e,f) was better at predicting water uptake in each soil layer due to its more realistic treatment of the dissipation of water potential gradients across layers. Accordingly, the model performed best out of the three at predicting the time-course of soil moisture (2.37% RMSE overall, Fig. 2, right). It did have trouble simulating nocturnal outflows at 50cm (Fig. 3g,h), leading to a slight dry bias at this depth over time. This was matched by a slight wet bias in the two deepest layers (100 and 150cm), which may be the source of hydraulic redistribution to 50cm in the data. In all layers except at 50cm, RMSE of soil predicted moistures was below 0.5%, in all other layers with a diurnal uptake signal model fit (Nash Sutcliffe Efficiency) was above 0.75, and in all layers the overall model bias was below 0.05 mm/d, outperforming the Ohm’s law models (Fig. 4). By better representing water potential gradient dissipation than either Ohm’s law model, the big root model was able to significantly reduce biases in water uptake and soil moisture predictions.

**Figure 4:**
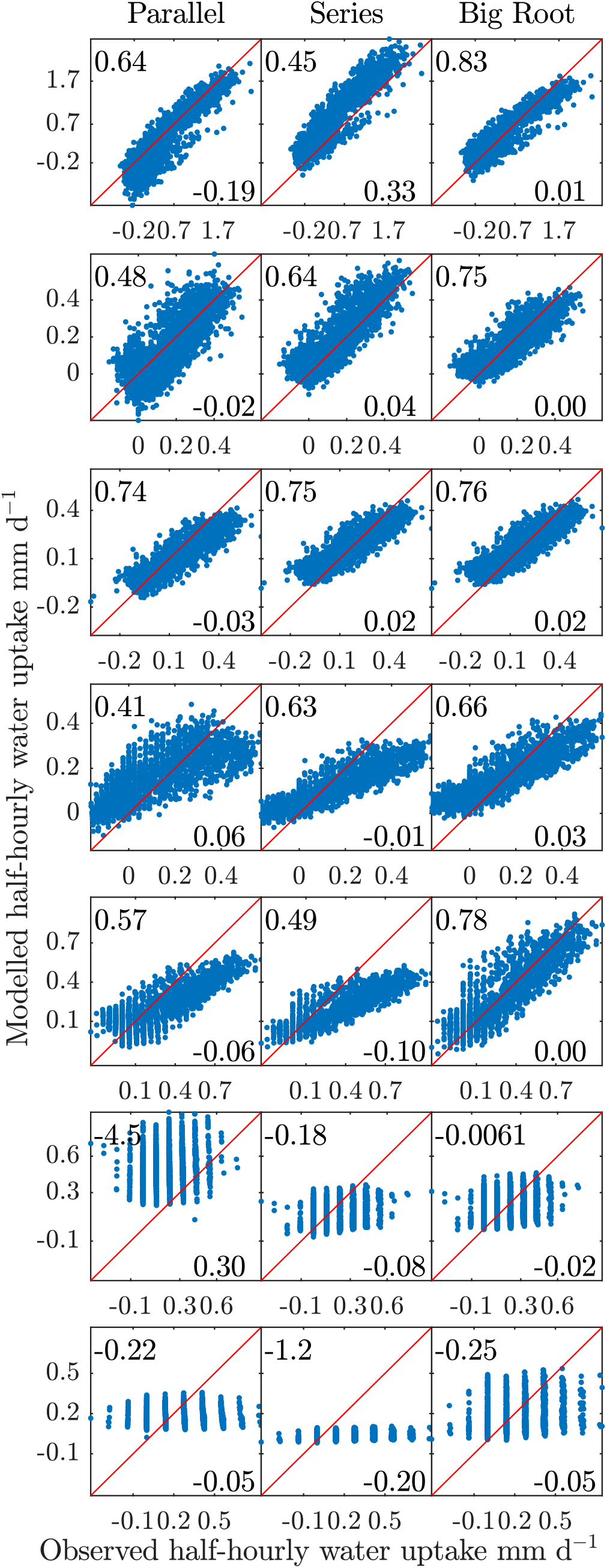
Model vs. observed half-hourly water uptake (mm/d) covering entire period, for each model (columns) and all dephts (rows); 1:1 line in red, Nash-Sutcliffe Efficiency at top left and model bias (mm/d) at bottom right.

All three models had trouble with the dynamics of the two deepest layers (Fig. 3k-n), where observations did not exhibit diurnal cycling of moisture loss. This likely breached an assumption of the model calibration procedure (see Methods), that all water movement is through the vegetation. Of the three models, the big root model had the lowest bias in these layers, showing that while it did not simulate the diurnal course of uptake well given that assumption, it was best at predicting its mean level.

## Discussion

The trade-off in the big root model between water uptake in a given layer and the pressure difference across it approximates the effects of RSA in real root systems. Any Ohm’s law model must assume a specific RSA, rigidly imposing water potential gradient configurations that can lead to biases in water uptake and predicted soil moisture, as seen in the present results. Overpredicting hydraulic redistrbution, as the parallel RSA does, has the potential to overestimate total plant-available water over time, by always refilling drier layers. Underpredicting access to deep water, as the serial RSA does, has the opposite effect. These errors can thus propagate to soil temperatures or transpiration and related fluxes at greater time scales. Starting from the porous pipe model (Eq. 1), it is possible instead to derive analytically the representation of a single, effective root, whose hydraulic architecture mimics the RSA at a given site. By better tracking the dissipation of water potential gradients due to water uptake, the resulting ‘big root’ model was able to account for the complex effects of real RSA, leading to a significant reduction of soil moisture and water uptake biases. Accounting for the effects of RSA in water uptake is thus a step towards reducing important soil moisture, as well as transpiration and heat flux biases identified in CMIP5 simulations [8].

The physical basis behind the big root model supports the robustness of its predictions: it is constrained to always behave as a single (effective) root with a given set of hydraulic properties. Its mathematical form, eq. 2, reveals the model to be a special case of the RSA Stencil, a model that accurately predicts uptake by root systems of a variety of RSA [37]. The more general, unconstrained RSA Stencil can be made to fit diurnal uptake dynamics better than the constrained big root model, including nocturnal redistribution to the 50cm layer, but simultaneously fails to predict longer-term soil moistures (SI Numerical Results). The big root constraints on the RSA Stencil derived here thus sacrifice some of the flexibility of RSA Stencils in representing diverse RSA to achieve greater physical robustness. The robustness of predictions, based on accurate representations of mechanisms, is especially important in the context of future climate projections, where vegetation water uptake potentially takes place in no-analogue scenarios.

Given its physical basis, it is possible to use the model to find meaningful hydraulic parameters of the ‘effective root’ (i.e., total within-layer soil-to-root conductance and the cross-layer root conductance, see SI Big Root Model) rather than just constrained RSA Stencil coefficients that predict water uptake, as was done here. The necessary condition is to have data on xylem water potential at the base of the root system for each time-step, which can be obtained through stem psychrometry [44]. Obtaining time-series of how such effective ‘big root’ parameters change at different time scales in real vegetation may lead to insights concerning the importance of underlying biological mechanisms (e.g. root growth and senescence, aquaporin activity) at larger scales.

To allow for borader application of a big root model in LSMs, its parameter values need to be established for a number of sites with distinct plant functional types (PFTs). The calibration procedure is data-intensive and subject to some restrictive assumptions. It is particularly important to arrive at an adequate partitioning of root water uptake from soil water movement prior to calibration. Model calibration is also sensitive to the soil moisture characteristic (*θ-ψ_s_*) curve, as the model is linear in soil water potential, which is itself very nonlinear in soil moisture. Ideally, the calibration dataset would contain both sets of values, directly observed.

The main limitation of the big root model follows from the assumption that soil moisture is uniform in each layer. Lateral soil moisture heterogeneity is a key feature of unsaturated zone hydrology and neglecting it necessarily leads to model prediction error [34]. This assumption is not novel in the big root model, however, but is instead a feature of all LSMs. In fact, it was assumed at the outset of model development to ensure compatibility of the result with relevant LSMs and maintain sufficiently low computational cost. The model might, however, be extended to account for lateral soil moisture heterogeneity, for example by separately modelling a ‘dry’ and ‘wet’ soil column, with a two-root model simulating a split root situation. Micro-scale heterogeneity, between the rhizospehere and ‘bulk soil’ can also be the key limiting factor to soil-root flows [45]. While it was not represented in this study, this resistance can be modelled separately, in series with the RSA model [46]. Also, it is notable that not representing this mechanism in the big root model seemingly did little to degrade the quality of its predictions for the Wind River Crane data.

A big root model for site-scale vegetation water uptake can be derived from first principles describing water movement through plant roots. This formalism provides a simple and mechanistically based framework to describe a key process limiting land surface exchanges of matter and energy. Considering the non-linearity of potential gradients on this effective root performs better at predicting water uptake than commonly used Ohm’s law analogy models. Potential applications of the theory beyond terrestrial modelling include improving drought stress formulations in existing crop models, which often rely on empirical or heuristic formalisms to predict crop yields, and thus food production, in future climate scenarios.

## Methods

### Model Development

Solutions to eq. 1 can be used to define a mean value of water potential effective in root uptake,

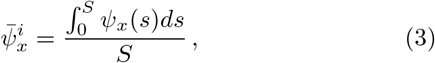

for each segment of a root of known length *S* and with uniform axial (*K_x_*) and radial (*k_r_*) hydraulic conductances. This is the value of water potential that allows uptake of the root segment to be found as 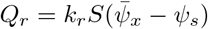. Solutions for this value,

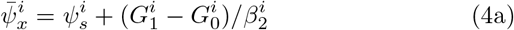

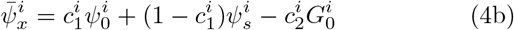

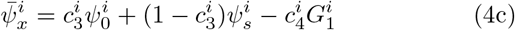

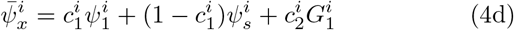

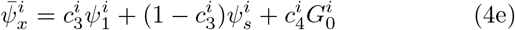

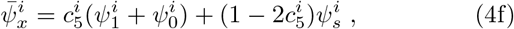

are linear in the layer boundary conditions: soil water potential 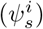 and the root water potential (*ψ^i^*) or its gradient (*G^i^*) at layer boundaries (subscript 0 at layer bottom and 1 at top); linear coefficients 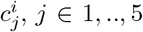 are determined by root properties 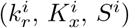, as shown in table 1.

**Table 1:**
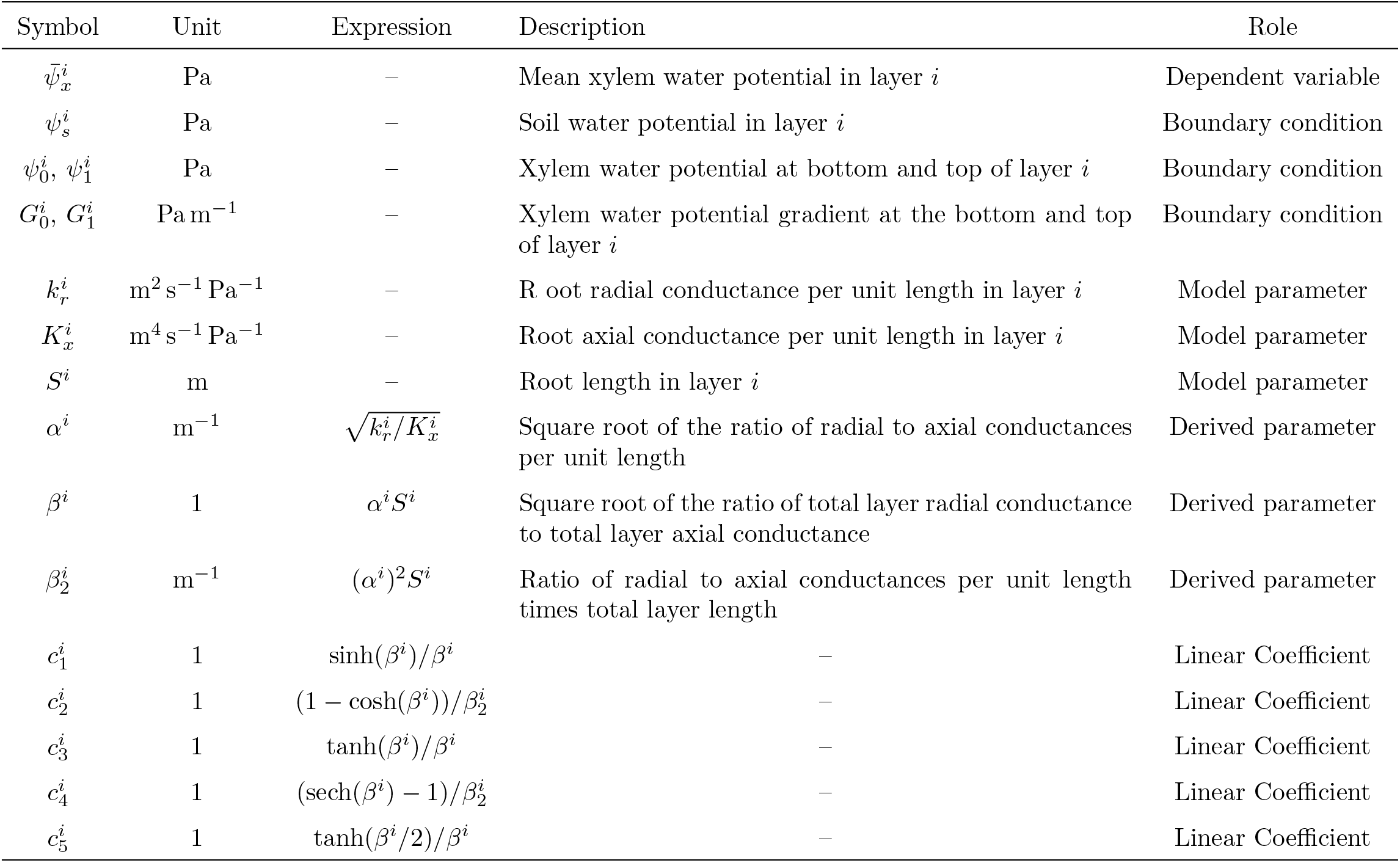
List of variables

Assuming a single root segment traversing each soil layer, these linear relations for 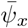 of adjacent layers can be combined and the conditions at layer boundaries cancelled (Fig. 5) to yield eq. 2. The linear coefficients in these equations are known functions of the root segment properties *k_r_, K_x_*, and *S* (further details of the derivation and resulting coefficient expressions in SI Big Root Model).

**Figure 5:**
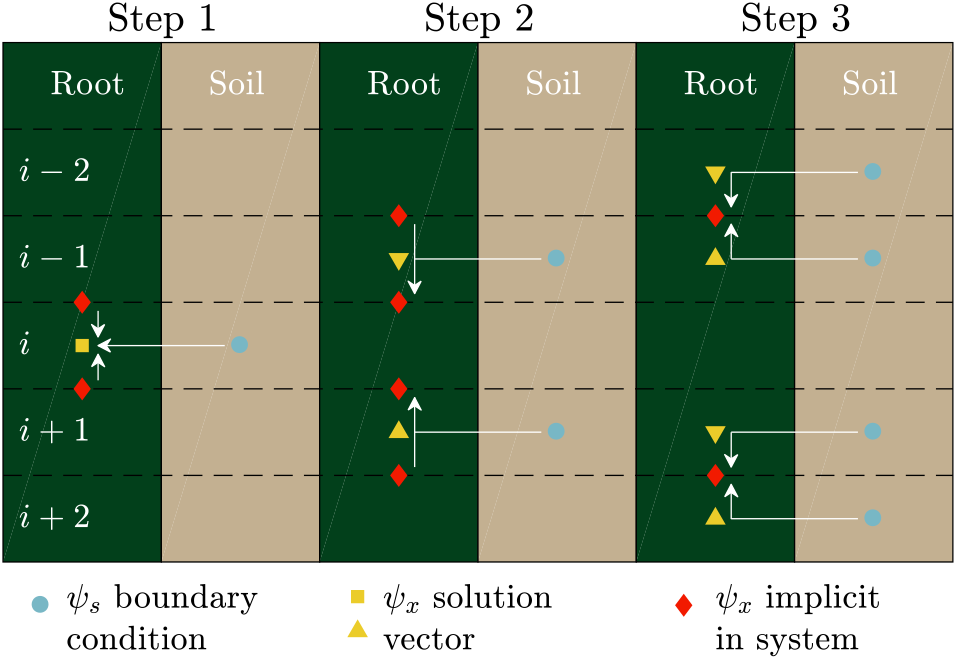
Assembly of big root linear equation for layer *i*. Step 1 selects an equation for 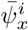 in boundary conditions at the top and bottom of the layer. Step 2 substitutes expressions for boundary conditions in layer *i* in boundary conditions at layers *i* ± 1. Step 3 replaces boundary conditions at layers *i* ± 1 with expressions in mean root and soil water potentials in layers *i* ± 1 and *i* ± 2. The result is a linear equation for mean root water potential in a given layer in terms of those in the four surrounding layers and the corresponding soil water potentials, which can be formulated as eq. 2.

Given a root of known properties, soil water potential, and boundary condition on water potential or its gradient at the stem base, a linear system constructed from eq. 2 may be solved simultaneously for the exact, analytical values of 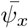 in all soil layers, also yielding the uptakes *Q_r_*. For a root of unknown properties on each segment, the model parameters *k_r_, K_x_*, and *S* can be estimated by inversion. By using the analytical relations between these parameters and the linear coefficients in eq. 2 as constraints when fitting the latter for unknown RSA, one calibrates a ‘big root’ model, which assumes that this RSA can effectively be represented as a single root.

### Site and Data

Wind River crane precipitation and soil moisture data at half-hourly resolution for the year 2010 [43] were used in conjunction with published fine root surface area profiles (Fig. 6a) and soil textures to define soil characteristic curve parameters [47, 48, 42]. The summer drought period was subselected from the data (Fig. 6b) as the period with no precipitation events that noticeably affect shallow-layer soil moisture. This was done in order to minimise soil water movement in the resulting dataset. Under the simplest assumption that all water movement during this period is due to plant water uptake or loss, root water uptake at each time-step was calculated from volumetric water content using centred differences. A calibration dataset of three 72-hour periods was further subselected from the drought period, at its start, middle, and end (Fig. 6c).

**Figure 6:**
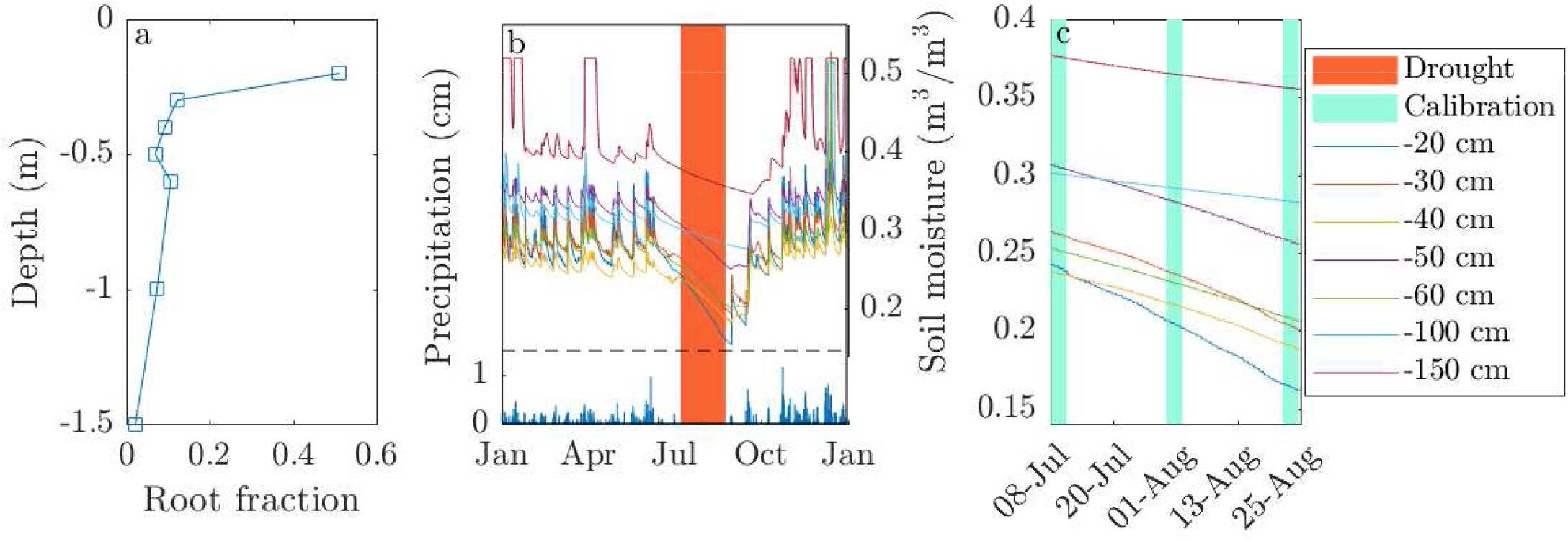
Wind River Crane data used: root profile (a) precipitation and soil moistures with drought period selected for simulation marked in orange (b). Detail of soil moisture at 7 depths during the selected drought period (c), with calibration data subset marked in cyan.

### Numerical Simulations

Calibration data included soil water potentials and calculated root water uptakes in each layer, as well as a flow boundary condition, found as the sum of water losses across all layers. The time-course of volumetric water content was not included in calibration data. The big root model was calibrated on the subset of three 72-hour periods. The inversion procedure estimated model parameters *k_r_, K_x_*, and *S* in each layer, although the absolute parameter values themselves are meaningless given the boundary conditions used, and served only to yield correctly constrained linear coefficients for eq. 2, capable of predicting root water uptakes (see SI Big Root Model for details). Root conductances between layers in the Ohm’s law in-series model was calibrated by inversion to the whole drought period, as parameters of satisfactory quality could not be found with the calibration subset alone. All inversions were performed using the Levenberg-Marquardt algorithm [49, 50]. Conductances from the stem base to each soil layer in the Ohm’s law parallel model, as well as soil-root conductances in the in-series model, were set in proportion to the observed rooting profile.

Forward simulations with each model used soil moisture in each layer at the start of the drought period as the initial condition and solved for soil water contents at each subsequent time-step, with water potentials calculated from predicted soil moistures. The simulation employed a Crank-Nicolson time-stepping scheme and represented the *θ* and *ψ_s_* relationship implicitly, using an iterative solver to account for its nonlinearity [51, 52].

All computational work was done in Matlab, release R2018a [53]; all original code is available on request from the author.

## Acknowledgments

I am deeply grateful to Sonia Wharton and the team at Wind River Crane who collected and kindly provided the data underlying the test case. I gratefully acknowledge my colleagues, whose comments improved this manuscript. This work was supported by the Ministry of Education, Youth and Sports (Czech Republic) through the institutional project MSMT CZ.02.1.01/0.0/0.0/16 019/0000803 and the Large Infrastructures for Research, Experimental Development and Innovations project “IT4Innovations National Supercomputing Center LM2015070”.

