## Supplemental Information for "Big root approximation of site-scale vegetation water uptake"

Martin Bouda[1]

[1]Faculty of Forestry and Wood Sciences, Czech University of Life Sciences in Prague, Kamýcká 129, 165 21, Prague, Czech Republic

#### Big Root Model

##### Model assumptions and problem setup

Root water uptake is governed by the 'porous pipe' equation,

$$\frac{d^2\psi_x}{ds^2} = \frac{k_r}{K_x}(\psi_x - \psi_s) \tag{S1}$$

where $K_x$, $k_r$ are root axial and radial conductances, respectively, $s$ is length along root, $\psi_x$ is unknown root water potential, and $\psi_s$ is the soil water potential [1]. An analytical solution of this equation for a single root with homogeneous properties [1] has recently been extended to root segments with properties varying in according to specific functional forms [2] and root systems of generalised hydraulic architecture [3, 4].

Terrestrial models divide the subsurface into $n$ depth layers, which are assumed to have homogeneous hydraulic properties. Soil hydrological equations, describing the movement of water in porous media, are discretised over these layers to obtain solutions for volumetric water content $\theta$ and water potential $\psi_s$ in each layer. Water uptake is calculated in line with the resulting hydration status of each soil layer and the evaporative demand given by atmospheric conditions and a model for photosynthesis and stomatal transpiration. So as to fit neatly with such a description of subsurface hydrology, this work assumes that both the plant and the soil are subdivided into depth layers with homogeneous properties. Aside from a single value of $\psi_s^i$ in each soil layer, the big root model also assumes a single value of the conductances $K_x^i$ and $k_r^i$ for the root and seeks to solve a system of equations for a single value of root xylem water potential, $\bar{\psi}_x^i$, in each layer, $i \in 1, .., n$, which represents the value effecitve in root water uptake, i.e. the value that correctly yields water uptake $Q_r$, when used in the equation $Q_r = k_r S(\bar{\psi}_x - \psi_s)$.

If we assume that each layer is traversed by a single root, we can use the definition of the mean of a continuous function to write an expression for $\bar{\psi}_x^i$ (eq. 3). The integral in this expression can be evaluated building on previous work on single [1] and multiple [3, 4] root solutions, to yield eqs. 4a-f.

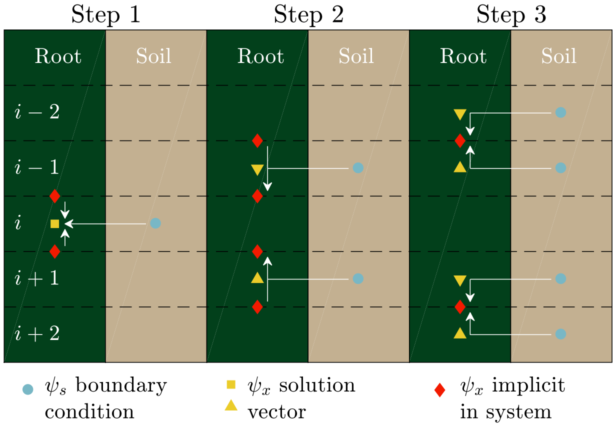


Figure S1: Assembly of big root linear equation for layer $i$. Step 1 selects an equation for $\bar{\psi}_x^i$ in boundary conditions at the top and bottom of the layer. Step 2 substitutes expressions for boundary conditions in layer $i$ in boundary conditions at layers $i\pm1$. Step 3 replaces boundary conditions at layers $i\pm1$ with expressions in mean root and soil water potentials in layers $i\pm1$ and $i\pm2$. The result is an equation for mean root water potential in a given layer in terms of those in the four surrounding layers and the corresponding soil water potentials.

##### Assembly of linear systems

Systems of linear equations assembled from the relations 4a-f can be used to find simultaneous solutions for $\bar{\psi}_x^i$ in all layers $i \in 1, .., n$. The assembly of such systems can be illustrated in three steps (fig. S1). The assembly of a system using layer boundary water potential gradients ($G_0^i$, $G_1^i$) as boundary conditions begins (Step 1) from eq. 4a, where the boundary conditions are as yet unknown. To eliminate the unknown $G_1^i$ (Step 2), substitute from eq. 4a evaluated in layer $i-1$ (note that $K_x^i G_1^i = K_x^{i-1} G_0^{i-1}$ from conservation of mass). The equation now contains the unknown $G_1^{i-1}$. An expression for this boundary condition can be found (Step 3) by combining eq. 4b evaluated in layer $i-2$ with eq. 4d in layer $i-1$. As $\psi_0^{i-1} = psi_1^i$ (from continuity), we can solve both 4b and 4d for the water potential at the boundary between the two layers and set the

Table S1: Expressions for coefficients in eq. S2 for different choices of implicit boundary conditions in assembly

| Definitions | | | Assembly using gradients at layer interfaces | | Assembly using potentials at layer interfaces | |
|---|---|---|---|---|---|---|
| Symbol | Expression | Layer | $\xi$ | $\sigma$ | $\xi$ | $\sigma$ |
| $r_K^{j,k}$ | $K_x^j/K_x^k$ | $i-2$ | $\frac{r_K^{i-1,i} r_{c1}^{i-2,i-1}}{\beta_2^i C_G^{i-2,i-1}}$ | $\frac{r_k^{i-1,i} r_{c1}^{i-1,i-2}(1-c_1^{i-2})}{\beta_2^i C_G^{i-2,i-1}}$ | $\frac{c_5^i}{c_2^{i-2} r_K^{i-1,i-2} C_\psi^{i-1,i-2}}$ | $\frac{c_5^i(1-c_1^{i-2})}{c_2^{i-2} r_K^{i-2,i-1} C_\psi^{i-2,i-1}}$ |
| $r_{cl}^{j,k}$ | $c_l^j/c_l^k$ | $i-1$ | $\frac{r_K^{i-1,i}}{r_B^{i-1,i} - \beta_2^i C_G^{i-2,i-1}}$ | $\frac{r_k^{i-1,i}(r_B^{i-1,i} - (1-c_1^{i-1}))}{\beta_2^i C_G^{i-2,i-1}}$ | $\frac{c_5^i}{c_2^{i-1} C_\psi^{i-2,i-1}} - r_{c5}^{i,i-1}$ | $\frac{c_5^i(3-c_1^{i-1}-1/c_5^{i-1})}{c_2^{i-1} C_\psi^{i-2,i-1}}$ |
| $r_B^{j,k}$ | $\beta_2^j/\beta_2^k$ | $i$ | – | 1 | – | $1-2c_5^i$ |
| $C_G^{j,k}$ | $c_2^j + c_2^k r_K^{j,k} r_{c1}^{j,k}$ | $i+1$ | $\frac{r_K^{i+1,i}}{r_B^{i+1,i} - \beta_2^i C_G^{i+2,i+1}}$ | $\frac{r_K^{i+1,i}(r_B^{i+1,i} - (1-c_1^{i+1}))}{\beta_2^i C_G^{i+1,i+2}}$ | $\frac{c_5^i}{c_2^{i+1} C_\psi^{i+1,i+2}} - r_{c5}^{i,i+1}$ | $\frac{c_5^i(3-c_1^{i+1}-1/c_5^{i+1})}{c_2^{i+1} C_\psi^{i+1,i+2}}$ |
| $C_\psi^{j,k}$ | $c_1^j/c_2^j + r_K^{j,k} c_1^k/c_2^k$ | $i+2$ | $\frac{r_K^{i+1,i} r_{c1}^{i+1,i+2}}{\beta_2^i C_G^{i+1,i+2}}$ | $\frac{r_K^{i+1,i} r_{c1}^{i+1,i+2}(1-c_1^{i+2})}{\beta_2^i C_G^{i+1,i+2}}$ | $\frac{c_5^i}{c_2^{i+2} r_K^{i+1,i+2} C_\psi^{i+1,i+2}}$ | $\frac{c_5^i(1-c_1^{i+2})}{c_2^{i+2} r_k^{i+1,i+2} C_\psi^{i+1,i+2}}$ |

resulting expressions equal to each other. Rearranging yields an expression for $G_1^{i-1}$, which can then be then substituted into the overall equation. Analogous steps are used to eliminate $G_0^i$, only using expressions evaluated in layers $i+1$ and $i+2$.

The resulting equation can be algebraically rearranged to the form

$$\bar{\psi}_x^i = \sum_{j=i-2}^{i+2} \xi^j \bar{\psi}_x^j + \sum_{k=i-2}^{i+2} \sigma^k \psi_s^k,\ (j \neq i), \tag{S2}$$

which shows that the mean xylem water potential in a given layer is a linear function of the mean xylem and soil water potentials spanning the two surrounding layers. This form of the equation also shows it to be a special case of the RSA Stencil model of [5]. Expressions for coefficients $\xi$ and $\sigma$ for each layer in this special case are given in table S1.

A toal of sixteen separate systems can be derived, by choosing different implicit boundary conditions. Complementary pairs of systems, used together, enable inversion of the model. The complement to the system described above is that, which uses $\psi_0^i$ and $\psi_1^i$ as boundary conditions in all layers instead of gradients. To derive this, we take eq. 4f as the basis in step 1 and again for the substitution in step 2. In step 3, we use eqs. 4b and 4d again, but eliminate the gradient terms to solve for values of $\psi_x$ at the boundary between layers $i-2$ and $i-1$ ($i+1$ and $i+2$). After rearrangement, we retrieve the form of eq. S2, albeit with new expressions for coefficients $\xi$ and $\sigma$ (again, given in table S1). Each of these separate parameterisations yields a distinct system of linear equations (with unequal parameters), which, however, yield the same solutions for a given set of boundary conditions.

#### Domain boundary conditions

The root is assumed to have no flow at the lower boundary, consistent with the 'porous pipe' model, simplifying the resulting equations for layers $n-1$ and $n$. This assumption follows from the idea that the rooting-depth is located somewhere in the last layer and thus no water uptake takes place deeper down. It may break down where vegetation has access to the phreatic zone, if this is deeper than the modelled soil domain. In such a case, an alternate boundary condition can be used.

Boundary conditions at the top of the domain may either specify the water potential ($\psi_C$) or gradient ($G_C$) at the stem base (or root collar). In either case, the fixed value replaces the respective expression for the boundary condition in the equations for layers 1 and 2, simplifying the resulting coefficients. For transient simulations of root system water uptake from a numerically represented soil, these values can vary at every time-step to simulate water demand from the above-ground portion of the vegetation. In the present study, these boundary condition forcings were derived from data, but for more sophisticated modelling, they could result from coupling with an above-ground plant and atmospheric model.

#### Simultaneous solution

A system of linear equations S2 for successive layers, with boundary conditions appropriately incorporated, can be rearranged into the form

$$\mathbf{Ax} = \mathbf{b}, \tag{S3}$$

where $\mathbf{A}$ is the matrix of coefficients $\xi$, $\mathbf{x}$ is the solution vector of values $\bar{\psi}_x^i$ for $i \in 1,..,n$ and $\mathbf{b}$ is the vector of values resulting from the second term on the right hand side of eq. S2 and boundary conditions in layers 1 and 2. Thus, if we know the root parameters $k_r$, $K_x$, and $S$ for each layer, then for any set of boundary conditions $\psi_C$ or $G_c$ and $\psi_s^i$ in each layer, this

Table S2: RMSE in converged big-root fits to stochastically generated synthetic root cases, mean values of root properties and solutions for comparison, aggregated across all 20 cases and all 7 layers.

| Data | Bdry. cond. | n converged | $\psi_x$ | $Q_R$ | $k_r S$ | $K_x/S$ |
|---|---|---|---|---|---|---|
| $\bar{\psi}_x$, $Q_R$ | $\psi_B$ | 18 | 3.9e-10 | 5.0e-19 | 3.1e-25 | 1.7e-24 |
| $\bar{\psi}_x$, $Q_R$ | $G_B$ | 19 | 3.8e-10 | 4.9e-19 | 3.4e-25 | 2.8e-24 |
| $Q_R$ | $\psi_B$ | 20 | 4.1e-10 | 4.6e-19 | 1.1e-24 | 4.8e-24 |
| | | | (2.4e4) | (6.3e-19) | (2.2e-10) | (5.4e-10) |
| $Q_R$ | $G_B$ | 20 | 1.2e4 | 4.7e-19 | 1.3e-10 | 3.0e-10 |
| actual values (mean ± st. dev.) | | 20 | -6.4±1.7e5 | 2.1±2.7e-4 | 1.4±0.6e-9 | 1.1±0.5e-9 |

pentadiagonal system provides the analytical, exact solution for the resulting $\bar{\psi}_x^i$ in each layer. This is termed here the 'big root' model.

#### Inverting the model

When the properties of the root are unknown, the linear coefficients can be estimated by inversion. To examine the quality of parameters estimated by standard model inversion methods, 20 cases of a single root with randomly varying properties ($k_r$,$K_x$,$S$) across 7 layers were generated. Central values ($L$=5 cm, $k_r$=2.83e-8, $K_x$=5.03e-10) were multiplied by a factor distributed uniformly between 0.5 and 1.5, independent in each layer. Values for $\bar{\psi}_x$ and $Q_R$ in all layers were then found analytically for each case for all combinations of the following boundary conditions: $\psi_B$ varied from -0.4 to -1.2 MPa and $\psi_s$ from -0.2 to -1MPa, each by increments of 0.2MPa. Big root stencis were then fit to these solutions using the soil water potentials as boundary conditions, as well as either the water potential at the stem base or its gradient.

It has already been shown that parameters $\xi$ and $\sigma$ are non-unique, as multiple formulas can be obtained analytically, which yield distinct values. While each parameter set, correctly composed into a linear system, is separately capable of producing exact values of $\bar{\psi}_x^i$, preliminary work showed that convergence rates are inadequate when using only a single set of parameter constraints (one or the other expressions for $\xi$ and $\sigma$ in table S1), leading to stalled optimisation runs or very high numbers of iterations (results not shown). By contrast, simultaneously using complementary sets of expressions (e.g. both sets in table S1) in the inversion leads to high rates of convergence (Table S2). Either of the two complementary sets of coefficients can be used in predictive runs, as they yield identical results.

The relation between linear coefficients ($\xi$,$\sigma$) and the underlying root parameters ($k_r$, $K_x$, $S$) is non-unique, as only ratios or products of the latter occur in the expressions for the former. Inversion of the model thus cannot yield these parameters directly. Exactly what parameters model inversion can constrain and how well the data are fit by the resulting parameter sets depends on the information used. Table S2 shows the errors in model fits and in the values of the product $k_r S$ (total layer soil-root conductance) and the ratio $K_x/S$ (integrated layer axial conductance) resulting from model inversions. Using data on both $\bar{\psi}_x^i$ and $Q_R^i$ for each layer to constrain the model, the ratios $k_r^i S^i$ and $K_x^i/S^i$ are correctly estimated, yielding correct values of $\beta^i$ (and thus linear coefficients) but not $\beta_2^i$ (though errors across layers offset to yield correct linear coefficients for both systems used in the inversion). This is true regardless of the boundary condition set at the stem base (water potential $\psi_B$ or its gradient $G_B$).

As field data are very unlikely to contain target values for $\bar{\psi}_x^i$, and the present site data do not, a second set of inversions used soil-root flows alone as the data. In this case, with a boundary condition on $\psi_B$, the same outputs are obtained as above, except in the $n^{\text{th}}$ layer. In the $n^{\text{th}}$ layer, a different set of linear coefficients is obtained that correctly predicts flows ($Q_R^i$), despite implying incorrect values of $\beta^n$. This shows that for the degenerate case that occurs at depth (because $G_0^n = 0$), the values of $k_r S$ and $K_x/S$ (and their combination, $\beta$) are not uniquely determined by the flow solution alone and the inversion converges on one of a set of equivalent stencils ($\xi$, $\sigma$). With a boundary condition on $G_B$, correct $\beta^i$ values are not retrieved in any layer, though again a stencil (set of linear coefficients $\xi$, $\sigma$) yielding the correct $Q_R$ values is found.

Depending on the data used in model inversion, different interpretive value thus can (or cannot) be ascribed to the 'effective' big-root parameters underlying resulting stencil. These results suggest that measuring stem base water potentials is a valuable component of field site design if physically meaningful big root parameters are sought.

Table S3: Computational cost of a single run of each model, showing cpu time in s of a single matrix inversion (flow calculation), the number of these evaluations in the entire run, and the resulting total time spent calculating flow by each model.

| Model | cpu time (s / eval.) | n evaluations | total cpu time (s) |
|---|---|---|---|
| Big root | 1.6559e-5 | 130928 | 2.168 |
| Series | 1.6706e-5 | 148208 | 2.4760 |
| Parallel | 7.3619e-6 | 150504 | 1.108 |

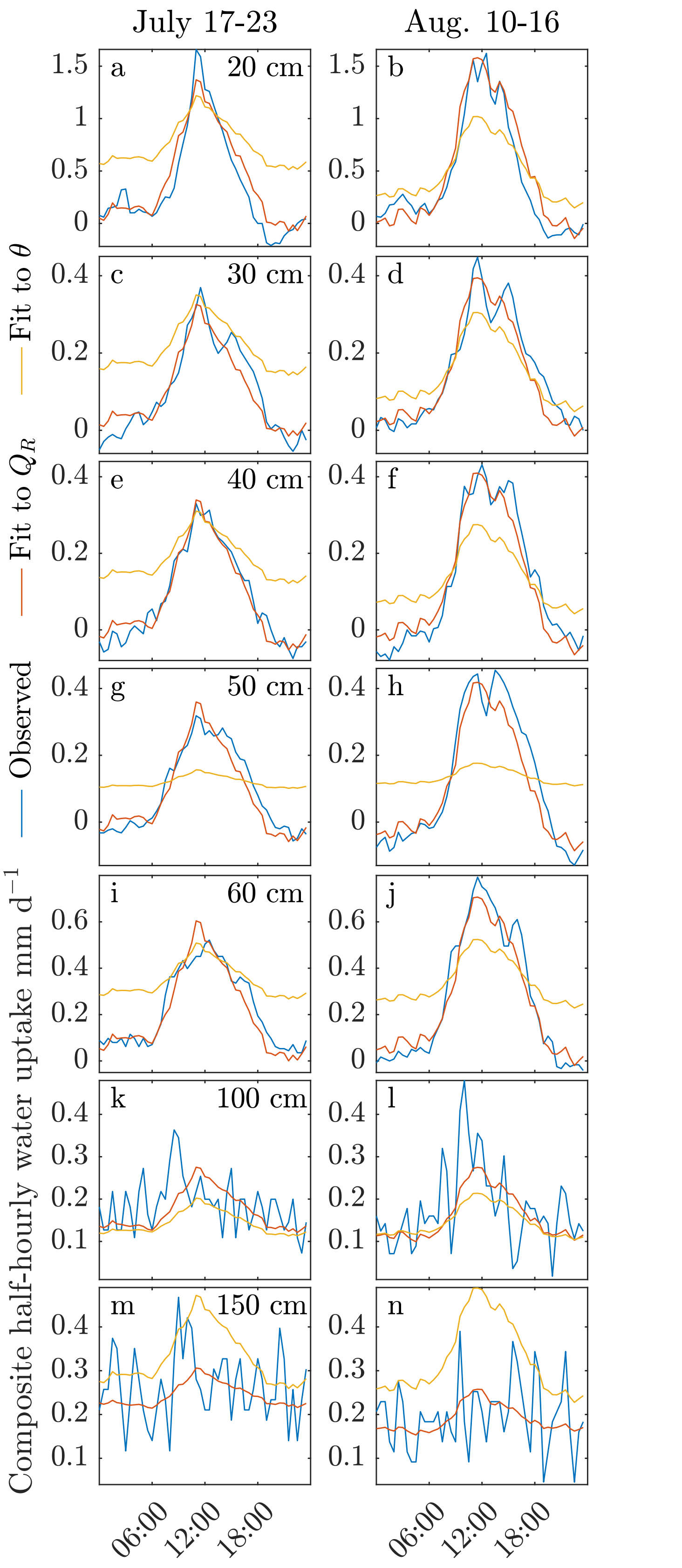


Figure S2: Comparison of observed and modelled half-hourly water uptake rate (mm/d) for 6-day composites of periods outside calibration dataset, displayed separately for each depth layer.

### Numerical results

#### Computational cost of Ohm's law and big root model runs

The computational cost of the prediction run with each of the models (Ohm's law series, parallel, and big root) is summarised in Table S3. While there were 2280 time steps in the simulation, the number of evaluations of each model are higher, as the nonlinear $\theta$-$\psi_s$ relationship was implicit in the equations solved, requiring an iterative solver. The number of evaluations of each model thus reflects the influence that the choice of root uptake model has on the convergence rate of the solution of the soil moisture characteristic curve. The big root model and resistances-in-series model had comparable cpu time cost per solution of the linear system for $\bar{\psi}_x$ and the difference in their total cost is attributable to the different convergence rates between them. The resistances-in-parallel model in the present implementation did not actually solve for $\bar{\psi}_x$, as this would involve an arbitrary partitioning of the total soil-stem base resistance into a soil-root and a root-stem base component. This is reflected in the lower cost achievable by this model. It should be noted, however, that implementations that vary soil-root and root-base resistances explicitly (e.g., due to xylem cavitation or rhizosphere characteristics) would face a per-evaluation cost about equal to the other two models.

#### Unconstrained RSA Stencil fits

RSA Stencils can be fit to the Wind River Crane 2010 drought data without 'big root' constraints, following the methods of [5]. One such stencil was fit to the calibration subset of $Q_R$ data, in the same way as the big root model. Another was instead fit to the time-course of soil moisture ($\theta$) in each layer, minimising the difference between predicted and observed $\theta$ over the entire drough period. The results of these inversions demonstrate the relative benefits of using the big root constraints. Whereas the big root parameter optimisation converged after 13 iterations, (1.9 seconds, 314 function evaluations), the comparable unconstrained fit to $Q_r$ data was not fully converged asfter 2500 iterations (over 145000 function evaluations, 15 minutes). The low rate of convergence of the unconstrained model is likely due at least in part to their non-uniqueness (demonstrated even for the single root case) and the complexity of the model.

The resulting parameters of the unconstrained stencil nevertheless showed better fit to the $Q_R$ data (Fig. S2), including an ability to simulate nocturnal redistribution into the 50cm layer. They did not, however, achieve very good predictions of the time-course of $\theta$ (Fig. S3, left), showing a wet bias across layers. The unconstrained fit to $\theta$ over the whole drought, by contrast, achieved near-perfect fits to these data (Fig.S3, right), but at the expense of poor fits to the diurnal dynamics of uptake (Fig. S2).

Imposing big root constraints on the RSA Stencil thus not only facilitated convergence, but also improves the robustness of the resulting parameters. While their fit to the data used in

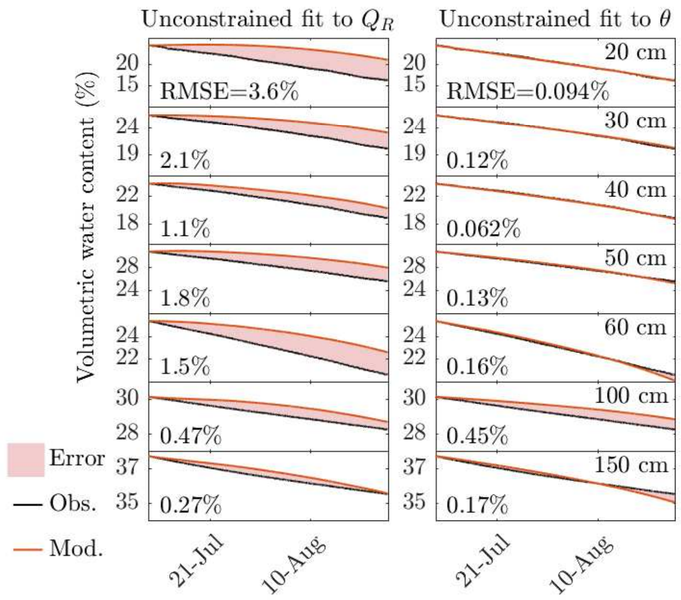


Figure S3: Observed (black) volumetric soil moisture (%) over the entire simulation period and values and predicted (red) by two unconstrained RSA stencils: one fit to uptakes (left) and the other to the soil moisture time series (right); results divided by soil layers (depths at right). Prediciton errors shaded in red. Root Mean Square Error of prediction reported at left of each panel.

the inversion is somewhat reduced compared to the unconstrained case, the physical consistency imposed by the big root assumptions improves the predicted flows in a way consistent with the overall time-course of soil moistures, even though they are not part of the calibration dataset. The constrained model is more physically robust and more easily yields physically consistent parameters than the unconstrained version. This also suggests a greater likelihood of the constrained model's robustness outside the range of calibration.
